# Virotrap Reveals *Salmonella* SopB as A Ubiquitinated Cargo for Host ESCRT-0

**DOI:** 10.1101/2025.08.19.669813

**Authors:** Margaux De Meyer, Annick Verhee, Hanna Grzesik, Delphine De Sutter, Jon Huyghe, Louis Delhaye, Tessa Van de Steene, Igor Fijalkowski, Veronique Jonckheere, Leander Meuris, Mathieu JM Bertrand, Petra Van Damme, Virginie Stévenin, Sven Eyckerman

## Abstract

The pathogenic bacterium *Salmonella* survives and replicates in host cells within a *Salmonella*-containing vacuole (SCV), a membranous niche it actively remodels via secreted effectors. The phosphoinositide phosphatase SopB is a key effector implicated in SCV biogenesis, yet its host protein interactome has remained incompletely defined. Using the mass spectrometry-based interactomics technology Virotrap, we identify a novel set of SopB-associated host proteins, including the ESCRT-0 subunit HGS and other membrane remodeling proteins. We demonstrate that SopB directly interacts with HGS via its ubiquitin-binding domains, and show that SopB promotes ESCRT-0 recruitment to the SCV, yet ultimately counteracts its anti-replicative effect. As the ESCRT machinery plays a central role in cargo sorting and membrane remodeling, we propose a new SopB-dependent strategy by which *Salmonella* hijacks host endosomal trafficking to build its intracellular niche.

## Introduction

*Salmonella* is one of the primary agents responsible for foodborne illnesses and diarrheal diseases globally [1] and is classified as a high-priority pathogen by the WHO [2]. *Salmonella enterica* subspecies *enterica* serovar Typhimurium (*S*. Typhimurium) typically induces self-limiting gastroenteritis in humans but can cause life-threatening systemic infections, particularly in immunocompromised individuals [3]. Infection is initiated by the active invasion of non-phagocytic intestinal epithelial cells through the induction of ruffles and bacterial uptake. Within enterocytes, the pathogen creates an intracellular niche known as the *Salmonella*-containing vacuole (SCV), which provides an environment that supports bacterial survival and replication [4]. In addition to its intravacuolar residence, other intracellular lifestyles are adopted by *S*. Typhimurium, with approximately 10% bacteria escaping the vacuole to hyper-replicate within the cytosol [5, 6], or persisting dormant salmonellae inside a divergent vacuole in human enterocytes [7]. The orchestration of invasion, SCV maturation, and intracellular pathogenesis is dictated by a repertoire of over 40 virulence factors, known as effectors, secreted through the type-3 secretion systems (T3SS)-1 and -2, encoded on *Salmonella* pathogenicity islands SPI-1 and -2, respectively [8, 9].

T3SS-1 effector SopB (*Salmonella* outer protein B), recently characterized as a phosphoinositide phosphotransferase [10, 11], has been implicated in a range of functions throughout the infection process [12, 13]. SopB contributes to efficient bacterial internalization by modulating phosphoinositide fluxes at the plasma membrane. These lipidic changes result in the activation of SH3-containing guanine nucleotide exchange factor (SGEF) that stimulate CDC42-dependent actin rearrangements, and promote membrane ruffling necessary for SCV and infection-associated macropinosome (IAM) formation [12, 14, 15]. Notably, ectopic expression of SopB in eukaryotic cells induces macropinosome formation [16]. SopB also promotes the acute synthesis of phosphatidylinositol (3,4)-bisphosphate (PI(3,4)P2) [10] thereby activating the host pro-survival kinase AKT that supports intracellular bacterial growth [17]. Shortly after cellular entry, SopB is mono-ubiquitinated at multiple lysine residues, which favors its translocation to the SCV and downregulates its activity at the plasma membrane [18, 19]. At the SCV, SopB orchestrates the early design of the intracellular niche by generating PI(3)P through recruitment of RAB5 and VPS34 [20]. This remodeling in membrane phospholipid composition enhances recruitment of sortin nexins SNX1 and SNX3, leading to spacious vacuole-associated tubules (SVAT) formation, SCV shrinkage and subsequent maturation [21–23].

Despite in-depth investigations into SopB’s phospholipid-modifying properties, ubiquitination-dependent localization [18, 19], and modulation of host cell signaling pathways [16], the links between these distinct, yet interdependent, biological aspects remain poorly understood. While the identification of protein interaction partners is commonly used for such mechanistic investigations, mapping of the full SopB interactome using classical approaches is hindered by SopB-specific challenges, including the high cytotoxicity resulting from SopB intracellular accumulation during overexpression and the potential requirement for a specific lipid environment [11]. An innovative interactomics strategy, called Virotrap, involves fusing a protein of interest to HIV-1 Gag to trap interacting protein complexes and identify them using mass spectrometry (MS) [24, 25]. By avoiding cell lysis, Virotrap enables a more comprehensive description of a protein’s micro-environment and interaction network, thus potentially revealing novel interaction partners. In this study, we leverage the unique properties associated with the Virotrap technology to characterize the SopB micro-environment and reveal a previously unidentified interaction between SopB and the host ESCRT machinery, which is typically involved in plasma membrane receptor sorting [26]. In particular, our findings reveal a direct, ubiquitin-dependent interaction between *Salmonella* SopB and the ESCRT-0 subunit HGS, both at IAMs and at the SCV. This newly identified interaction sheds light on how the extensively studied effector SopB a central endocytic host complex to manipulate membrane dynamics and support niche formation.

## Results

### Virotrap Charts the SopB Endocytic Membrane Environment

To gain new insight into the biology of the *Salmonella* effector SopB, we examined its host interactome using the Virotrap technology (**Fig. 1a**). This approach is based on the fusion of a protein of interest (the ‘bait’) with the human immunodeficiency virus-1 (HIV-1) Gag protein. Upon expression, Gag-bait fusions forms virus-like particles (VLPs) that encapsulate co-sorted interaction partners. Following purification of the released VLPs from the cell supernatant, mass spectrometry (MS)-based proteomics reveals the content of the particles, including interaction partners bound to the bait protein. This approach supports the characterization of weak and valency-dependent interactions within the bait micro-environment [27], including modification-dependent interactions [24, 28], and supports the analysis of proteins that may otherwise be cytotoxic when overexpressed [28]. Based on these properties, we generated a Gag-SopB fusion construct to enable Virotrap analysis. Western blotting confirmed that SopB fusion to Gag and expression in HEK293T cells resulted in SopB detection in VLPs collected from the culture medium (**Suppl. Fig. 1**). Next, we analyzed the proteomic content of Gag-SopB VLPs compared to a negative control VLP using MS, as previously described [24, 25]. In total, 478 human proteins showed differential abundance in VLPs containing SopB compared to the control (**Fig. 1b**). To investigate this extensive SopB interactome, we performed gene ontology enrichment analysis to identify associated cellular compartments (**Fig. 1c**). Consistent with SopB’s documented endosomal localization [29], Gag-SopB VLPs demonstrated significant enrichment of 47 endosomal proteins (**Fig. 1b, c**), including components of multiple endocytic host complexes (**Fig. 1c**). Intriguingly, all three subunits of the ESCRT-0 complex, namely HGS, STAM and STAM2, as well as the accessory ESCRT proteins HD-PTP and Endofin exhibited significant enrichment with the SopB bait protein (**Fig. 1b**). Together, these proteins form the machinery known to sort ubiquitinated cargoes, particularly membrane receptors, into intraluminal vesicles (ILVs) within multivesicular bodies [30].

**Figure 1.**
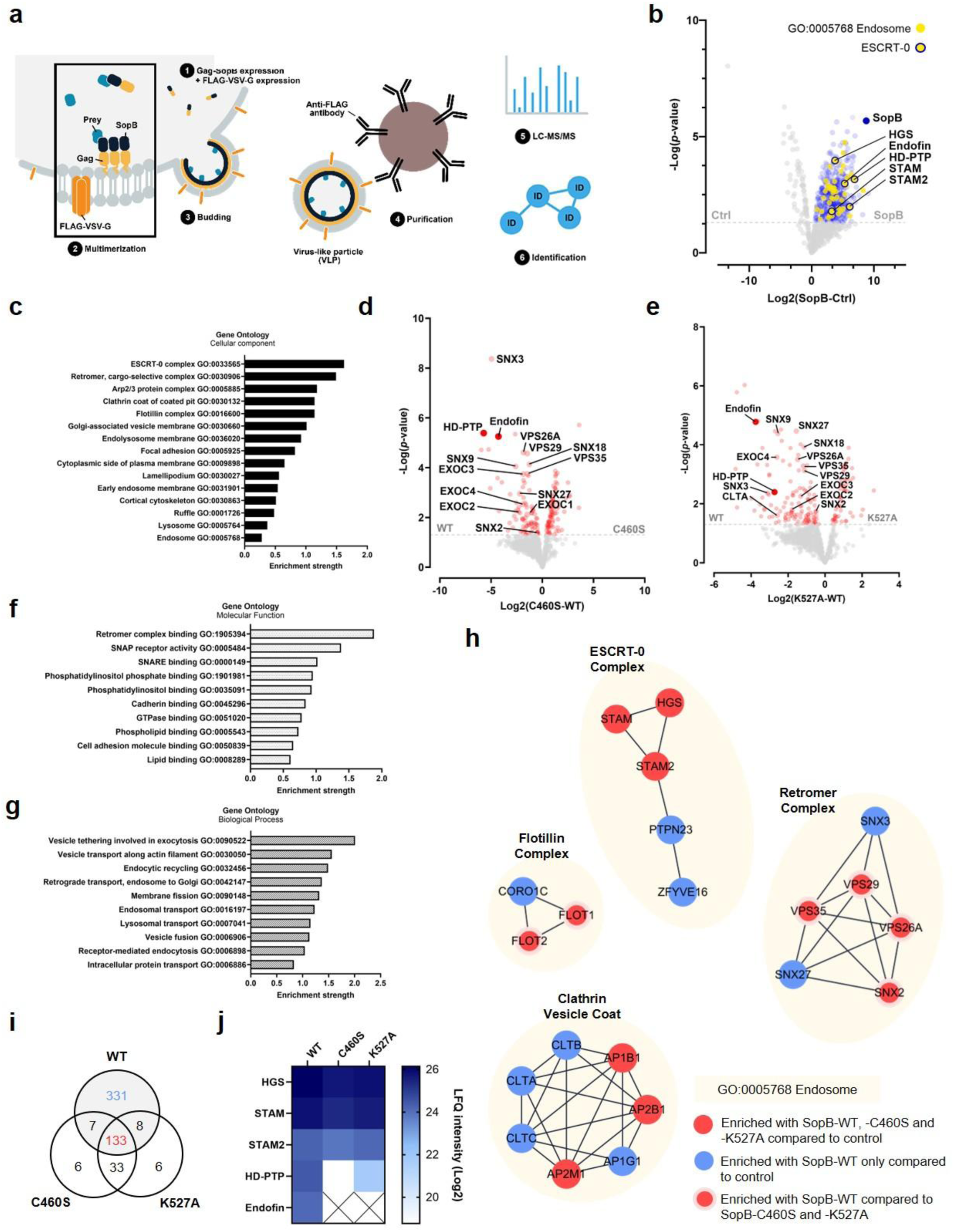
Virotrap maps the host environment of *Salmonella* SopB. (a) Overview of Virotrap. A fusion of the bait, e.g., effector protein of interest, to the C-terminus of the Gag polyprotein (myristoylated; pink zigzag line) of human immunodeficiency virus type 1 (HIV-1) is transiently expressed in HEK293T cells. Along with the Gag-bait fusion, FLAG-tagged vesicular stomatitis virus glycoprotein (VSV-G) is expressed and becomes embedded in the plasma membrane. Gag facilitates multimerization and localization of the fusion protein at the plasma membrane, giving rise to virus-like particle (VLP) budding. Consequently, host preys are trapped inside VLPs that can be captured in the cell medium by anti-FLAG immunoprecipitation. LC-MS/MS-based analysis of the VLP content subsequently allows identification of significant bait co-enriched host proteins. (b) Volcano plot displaying the differential enrichment between Gag-SopB wild-type (WT) and Gag-eDHFR in Virotrap (limma, p-value < 0.05). (c) Gene Ontology (Cellular Component) enrichment analysis for enriched proteins (p-value < 0.05) with SopB in Virotrap. STRING v12.0 was used for enrichment analysis. 15 significant terms are shown (FDR < 0.05). (d) Volcano plot displaying the differential enrichment between SopB-C460S catalytic mutant and SopB-WT in Virotrap (limma, p-value < 0.05). (e) Volcano plot displaying the differential enrichment between SopB-K527A catalytic mutant and SopB-WT in Virotrap (limma, p-value < 0.05). (f) Gene Ontology (Molecular Function) enrichment analysis for enriched proteins (p-value < 0.05) with SopB-WT compared to C460S in Virotrap. STRING v12.0 was used for enrichment analysis. 10 significant terms are shown (FDR < 0.05). (g) Gene Ontology (Molecular Function) enrichment analysis for enriched proteins (p-value < 0.05) with SopB-WT compared to C460S in Virotrap. STRING v12.0 was used for enrichment analysis. 10 significant terms are shown (FDR < 0.05). (h) Interaction network representation of significant endosomal protein complexes identified in Virotrap for SopB-WT. Connections (edges) depict high-confidence physical interactions with a STRING score of at least 0.7. (i) VENN diagram showing overlap of significant hits among all three SopB variant Virotrap screens upon comparison with Gag-eDHFR. (j) Heatmap representing LFQ intensities for ESCRT-0 components over all SopB variant Virotrap comparisons.

For a more in-depth functional characterization of the SopB interactome, we performed comparative Virotrap analyses using the catalytically inactive mutant SopB-C460S [31] and the catalytically reduced mutant SopB-K527A. Western blotting confirmed the presence of both mutants in VLPs collected from the culture medium (**Suppl. Fig 1**). This comparison revealed 86 significantly enriched hits for the WT effector form, including multiple sorting nexins (SNX2, SNX3, SNX9, SNX18, and SNX27), components of the cargo-selective retromer complex (VPS26A, VPS29, and VPS35), components of the exocyst complex (EXOC1/SEC3, EXOC2/SEC5, EXOC3/SEC6, and EXOC4/SEC8), clathrin chains (CLTA, CLTB, and CLTC) and ESCRT-associated proteins (HD-PTP and Endofin) (**Fig. 1d, e**). Highly similar results were obtained for the WT versus K527A comparison (**Fig. 1g**), indicating that reduced enzymatic activity impacts the SopB-host interactome. Gene ontology enrichment of the differential SopB interactome aligned with molecular functions and biological processes associated with SopB’s enzymatic activity, including “phosphatidylinositol binding” [11], “vesicle tethering in exocytosis” [32, 33], “retrograde transport, endosome to Golgi” [21], and “endocytic recycling” [34] among others (**Fig. 1f, g**). Across all Virotrap conditions, the SopB-WT, C460S, and K527A shared 131 significantly enriched proteins, while 333 hits were uniquely enriched in the SopB-WT profile (**Fig. 1i**), which underscores the importance of SopB catalytic activity in defining its interactome. Remarkably, accessory ESCRT-0 proteins HD-PTP and Endofin, highly enriched hits in the SopB-WT interaction profile, were not enriched with the catalytic mutants (**Fig. 1d, e, h, j**). In contrast, HGS, STAM and STAM2 were enriched independently of SopB catalytic activity, suggesting a two-step interaction: SopB first associates with ESCRT-0 core components, followed by the recruitment of accessory proteins dependent on its catalytic function. Taken together, these Virotrap results capture the SopB host microenvironment and its dependence on SopB’s enzymatic activity: notably pondering a novel interaction between the host ESCRT-0 complex and SopB.

### SopB Interacts with the Host ESCRT-0 Machinery

The detection of ESCRT-0 components HGS, STAM and STAM2 in both SopB WT and catalytic mutant interactomes suggests a catalytic activity-independent interaction. Therefore, we tested for potential direct interactions between these host factors and SopB, using a split-luciferase NanoBiT protein–protein interaction assay in HEK293T cells (**Fig. 2a**). This assay offers high sensitivity, making it well-suited for detecting transient or weak interactions in a cellular context [35]. HGS, STAM and STAM2 were genetically fused in both orientations to the NanoLuc C-terminal small fragment (SmBiT), whereas SopB was fused to the larger N-terminal fragment (LgBiT) (**Fig. 2b**). All possible fusion combinations were evaluated for luciferase complementation, measuring luminescence relative to the HaloTag negative control setup (**Fig. 2c, Suppl. Fig. 2**). We observed high relative luminescence for all ESCRT-0 proteins demonstrating direct interaction between SopB and these host proteins (**Fig. 2c**).

**Figure 2.**
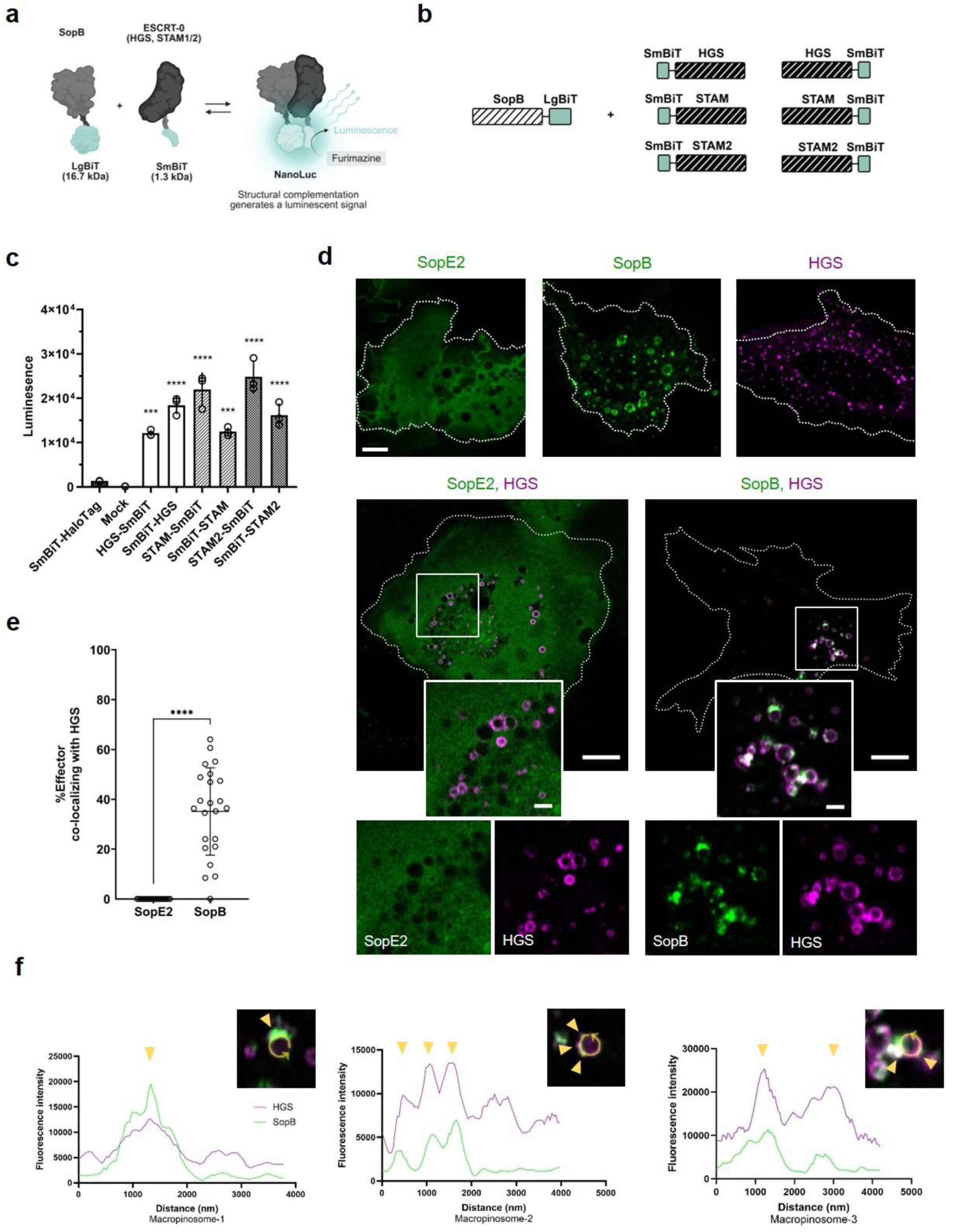
SopB interacts with host ESCRT-0 components. (a) Schematic representation of the NanoBiT protein-protein interaction assay. SopB (bait) and ESCRT-0 subunits (HGS, STAM, STAM2; prey) are genetically fused to either LgBiT- or SmBiT-encoding portions of the NanoLuc luciferase. Upon expression of the fusion proteins, the luciferase reconstitutes when an interaction between bait and prey occurs. After administration of the luciferase substrate furimazine, luminescence can be measured to detect a potential interaction. (b) Constructs used for NanoBit: LgBiT-tagged SopB with SmBiT-tagged HGS, STAM and STAM2 at either the N- or C-terminus. (c) NanoBiT protein-protein interaction assay evaluating the interaction between SopB C-terminally tagged with LgBiT and ESCRT-0 proteins HGS, STAM, and STAM2. The results represent three technical replicates. The experiment represents the results of three independent repeats. Statistical significance was determined using one-way ANOVA (***p < 0.001,***p-value < 0.0001). (d) Confocal imaging of stable inducible HeLa-HGS-mScarlet cells. Top row: SopE2-EGFP or SopB-EGFP transiently expressed without doxycycline. Middle and bottom rows: the same conditions with doxycycline induction (50 ng/mL); showing HGS-mScarlet localization relative to SopE2 or SopB). The scale bars indicate 10 µm (overview) and 2 µm (insets). Accessory ESCRT-0 proteins are indicated in non-transparent red. (e) Quantification of SopB and SopE2 co-localization with HGS. Data were collected from 26 cells from three independent experiments. Statistical significance was determined using an unpaired t-test (****p-value < 0.0001). Accessory ESCRT-0 proteins are indicated in non-transparent red. (f) Fluorescence intensity profiles of EGFP and mScarlet channels across the circumference of three representative macropinosomes. Arrowheads mark coincident peaks indicating SopB-HGS co-localization.

Next, we used live microscopy to assess the intracellular localization of SopB and HGS upon co-expression in epithelial cells. Heterologous expression of EGFP-SopB in HeLa cells revealed its localization to small intracellular vesicles, as well as macropinosomes formed upon SopB expression, consistent with previous studies (**Fig. 2d**) [29]. To monitor HGS localization with minimal overexpression artifacts, we engineered inducible HGS-mScarlet expressing HeLa cells allowing a controlled low HGS-mScarlet expression using an optimized low doxycycline concentration. We observed HGS-mScarlet localizing to small intracellular vesicles (**Fig. 2d**), which corresponds to its described endo- and lysosomal localization (Human Protein Atlas [36]). Upon SopB expression, HGS was also observed on SopB-induced macropinosome, where it co-localized with SopB. Further, to determine the role of SopB in HGS localization on macropinosomes, we monitored HGS localization upon SopB-independent macropinocytosis, by expressing the *Salmonella* effector SopE2 (**Fig. 2d**). Similar to SopB, SopE2 induced the macropinosome formation, and HGS was recruited to these structures, showing that HGS recruitment to macropinosomes does not depend on SopB. SopE2 displayed dispersed cytosolic localization whereas SopB showed a punctuate pattern localized to small intracellular vesicles and macropinosomes positive for HGS (**Fig. 2e**). Strikingly, SopB localized exclusively to HGS-positive areas. Especially, macropinosome membrane microdomains with high SopB fluorescence intensity co-localized with microdomains of high HGS fluorescence intensity (**Fig. 2f**). In summary, SopB co-localizes with HGS on macropinosome microdomains, but HGS recruitment to these structures is independent of SopB.

### HGS Localizes at *Salmonella* Infection Sites and Displays SopB-Dependent Recruitment at the SCV

Following the observed co-localization of ESCRT-0 and SopB on host endomembranes upon SopB heterologous expression, we next aimed to characterize the subcellular positioning of ESCRT-0 during infection in a SopB-dependent context. To this end, we performed time-lapse microscopy of stable inducible HeLa-HGS-mScarlet cells infected with fluorescent WT *Salmonella* (SL1344-WT) or SopB-deficient strain (SL1344-Δ*sopB*). Within the first five minutes after bacterial entry, we observed HGS recruitment to large vesicles and SCVs (**Fig. 3a**). Using fluorescent dextran, we confirmed that the large vesicles positive for HGS correspond to IAMs (**Fig. 3b**). Overall, HGS accumulation was observed on nearly all SCVs throughout the time-lapse (**Fig. 3c**) and on approximately 75 % of IAMs (**Fig. 3d**). Interestingly, infection with SL1344-Δ*sopB* led to a ∼25% reduction in HGS recruitment to SCVs (**Fig. 3c**), while recruitment to IAMs was not significantly affected (**Fig. 3d**). The latter observation is in line with the partial HGS-positive character of macropinosomes seen during SopE2 overexpression (**Fig. 2d**).

**Figure 3:**
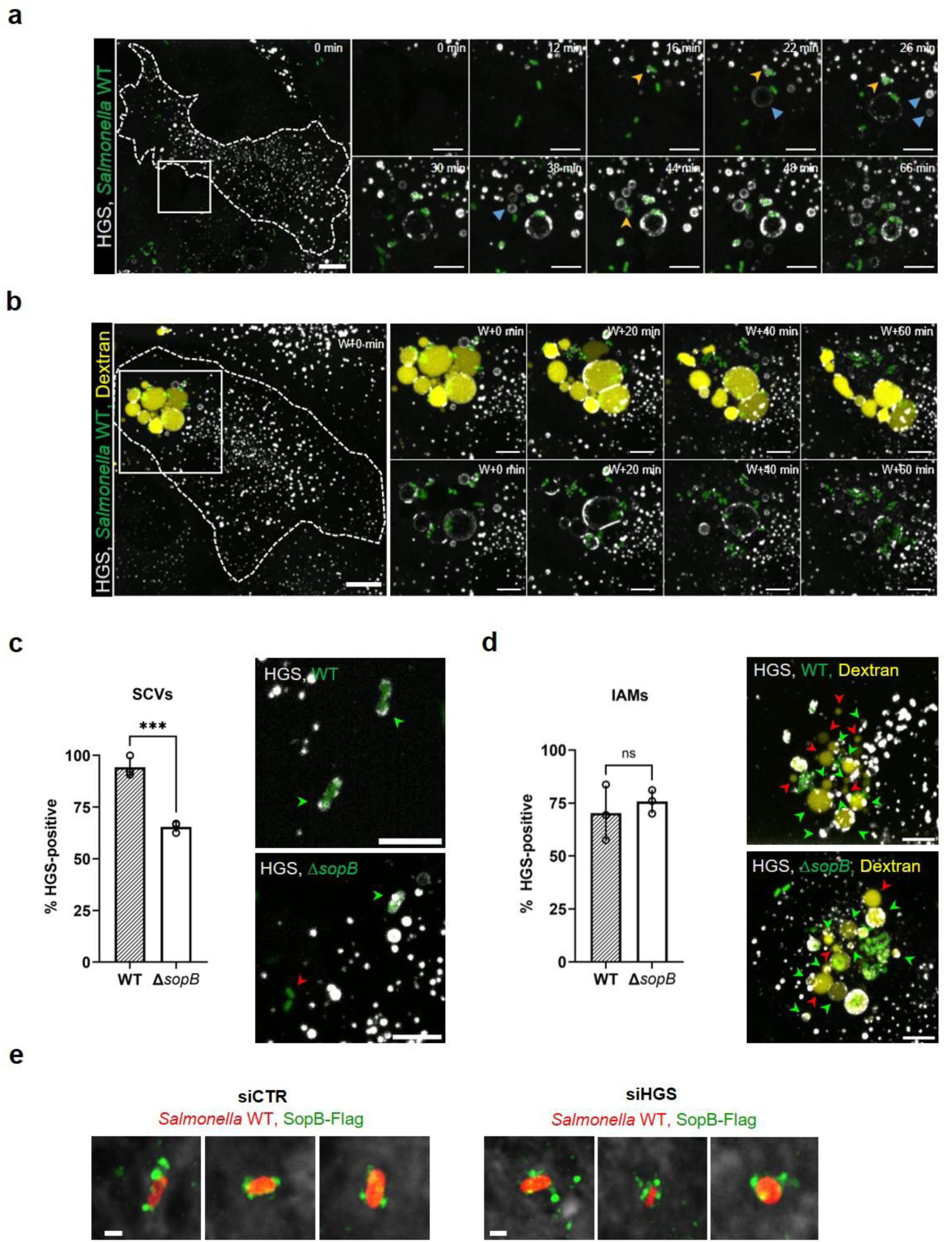
HGS is recruited at Salmonella infection sites. (a) Time-lapse imaging of HeLa cells expressing mScarlet-HGS upon doxycycline induction (white) and infected with GFP-expressing *Salmonella* SL1344 WT (in green). Bacteria are added to the cells just before the beginning of the acquisition to visualize entry and early events. See also **Suppl. Video 1** (cropped view). Yellow arrow-head: HGS-positive SCV, Blue arrow-head: HGS-positive IAMs. Scale bars: 10 µm (overview) and 5 µm (insets). (b) Time-lapse microscopy of HeLa cells expressing mScarlet-HGS upon doxycycline induction (in white) and infected with GFP-expressing *Salmonella* SL1344 WT (in green) in the presence of fluorescent Dextran (in yellow). Dextran and Bacteria are added to the cells for 30 min and washed (W) before the beginning of the acquisition to visualize IAM dynamics. See also **Suppl. Video 2** (cropped view). Scale bars: 10 µm (overview) and 5 µm (inset). (c) Quantification of SCVs that become HGS-positive over the course of the time-lapse in HeLa cells. Over 180 SCVs were scored across three independent experiments (mean ± SEM). Statistical significance was determined using an unpaired t-test (****p < 0.001). Arrowheads in representative images indicate HGS-positive (green) and HGS-negative (red) SCVs. The scale bars indicates 10 µm. (d) Quantification of IAMs that become HGS-positive over the course of the time-lapse in HeLa cells. Over 180 IAMs were scored across three independent experiments (mean ± SEM). Statistical significance was determined using an unpaired t-test (ns, not significant). Arrowheads in representative images indicate HGS-positive (green) and HGS-negative (red) IAMs. Scale bars indicates 5 µm. (e) SopB localization during infection using SL1344(SopB-Flag)-dsRed upon depletion of HGS. HeLa cells were infected and fixed at 30 minutes post-infection. SopB-Flag was visualized using mouse anti-Flag antibodies and donkey anti-mouse DyLight-488. Cells were stained using Deep Red cell mask (white). Scale bar indicates 1 µm.

Next, we examined SopB recruitment to the SCV with and without depletion of HGS with siRNAs. First, we monitored SCV and IAM formation upon HGS silencing by time-lapse microscopy using the PI(3)P biosensor 2xFYVE [37], and observed no significant differences in SCV or IAM formation compared to control cells (**Suppl. Fig. 2**). Subsequently, we infected the HGS-silenced HeLa cells with a *Salmonella* strain expressing a FLAG-tagged SopB followed by an anti-FLAG immunodetection. SopB localization to the SCV was not altered upon HGS depletion (**Fig. 3e**). Altogether, these data indicate that ESCRT-0 localizes to IAMs and SCVs—compartments known to associate with SopB during infection [23]—and support a model in which SopB promotes HGS recruitment to the SCV, rather than the reverse.

### SopB is Recognized as a Ubiquitinated Cargo for ESCRT-0

The ESCRT-0 complex (HGS, STAM and STAM2) directly binds ubiquitin moieties on cargo proteins destined for incorporation into ILVs within multivesicular bodies [38]. Taking advantage of the Virotrap MS data, we searched for diGly modifications indicative of ubiquitination sites and confirmed previously reported ubiquitination of SopB at lysines: K19, K41, K93 and K541 [18, 19, 39] (**Fig. 4a**). Notably, these ubiquitinations appeared independent of SopB’s catalytic activity in our Virotrap assay. Considering the physiological relevance of SopB ubiquitination of *in vivo*, this finding confirms the validity of using heterologous SopB expression to study interactions dependent on this post-translational modification. To assess whether interaction of SopB with ESCRT-0 is ubiquitin-dependent, four HGS domain deletion mutants were constructed, including removal of the VHS (Vps-27, Hrs and STAM) ubiquitin-binding domain, ubiquitin-interacting motif (UIM) domain, and a proline-rich region (PR) known to facilitate cargo binding independent of ubiquitination (**Fig. 4b-c**). NanoBiT protein-protein interactions assays revealed a marked reduction inrelative luminescence for HGS lacking ubiquitin-binding domains (**Fig. 4d**). While deletion of the VHS domain alone had little effect, ΔUIM led to a marked reduction in signal, identifying UIM as the main determinant for SopB binding. This confirms that SopB-HGS interaction depends on the ubiquitin-binding capacity of HGS.

**Figure 4.**
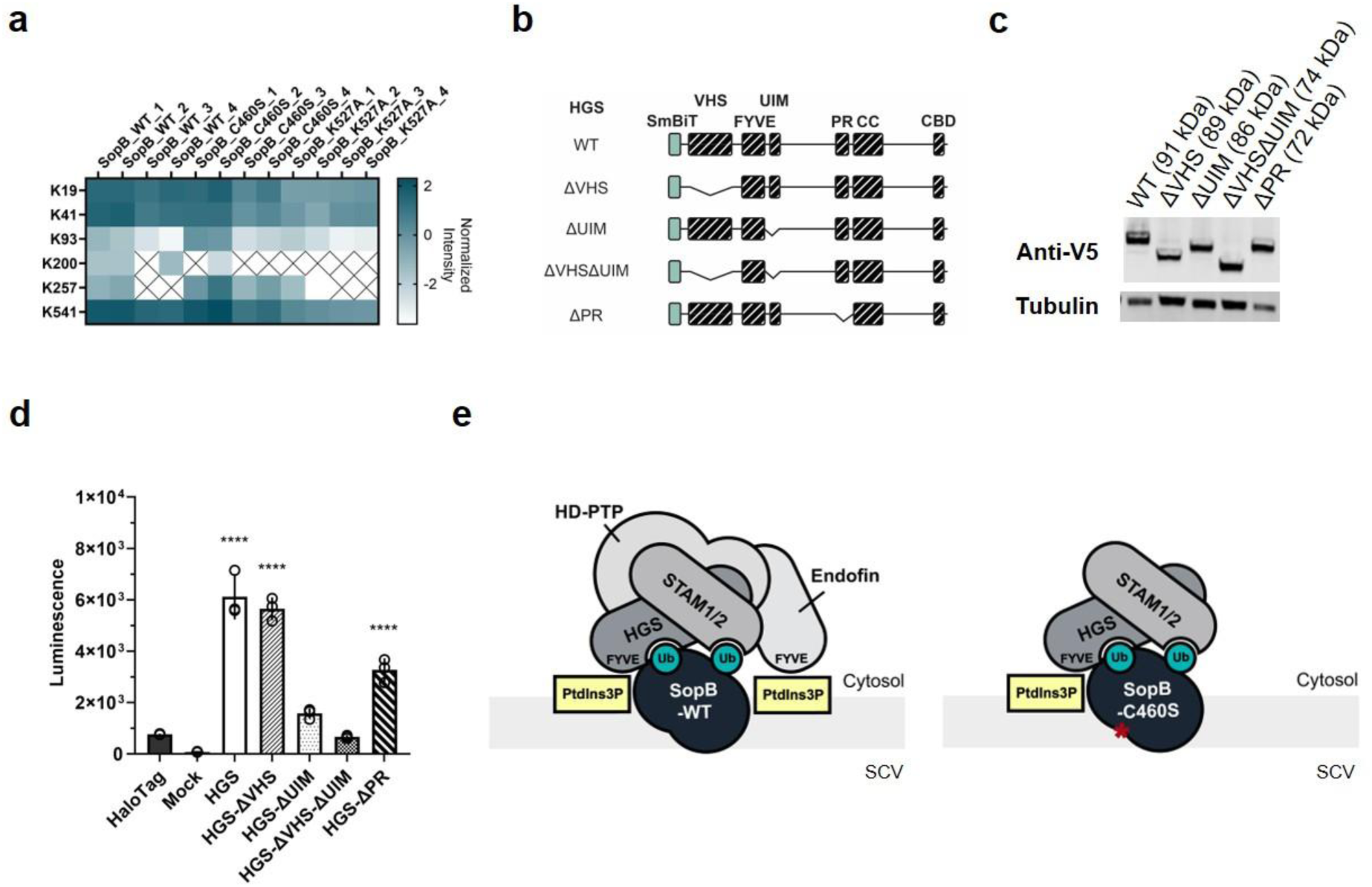
SopB is recognized as a ubiquitinated cargo for ESCRT-0. (a) Heat map of identified SopB peptides bearing diGly (K-ε-GG) remnants indicative of Lys ubiquitination across Virotrap datasets (WT and mutants). Intensities were normalized by median subtraction; crossed boxes indicate not detected. (b) HGS deletion constructs used in NanoBiT: VHS, Vps-27, Hrs and STAM; UIM, ubiquitin-interacting motif; CBD, clathrin-binding domain; PR, proline-rich; FYVE, Fab1p, YOTB, Vac1p, and EEA1 zinc finger. (c) Immunoblot analysis of SmBiT-V5-tagged HGS constructs expressed in HEK293T cells. Tubulin was used as a loading control. (d) NanoBiT assay measuring the interaction between SopB and HGS mutant variants. Statistical significance was determined using one-way ANOVA (*****p-value < 0.0001). (e) Model displaying ubiquitin-dependent binding of SopB with ESCRT-0 at the SCV and the differential recruitment of accessory host factors in comparison with a catalytic mutant SopB (Fig. 1).

In addition, deletion of the PR domain did not abolish SopB–HGS interaction, consistent with a proposed auxiliary ubiquitin-independent recognition, as previously described for interleukin (IL)-2 receptor recognition [40]. Since SopB ubiquitination was previously linked to its prolonged retention at the SCV [18], and given our observation that SopB promotes HGS recruitment to the SCV (**Fig. 3e**), we propose that SopB mono-ubiquitination serves as a molecular mimicry strategy to recruitHGS to the SCV (**Fig. 4e**). Following ESCRT-0 recruitment, the catalytic activity of SopB likely promotes subsequent recruitment of additional accessory proteins, as suggested by the comparative Virotrap analysis of WT and catalytic inactive SopB(**Fig. 1h, j**).

### ESCRT-0 Modulates *Salmonella* Infection Burden in Coordination with SopB

To investigate the functional relevance of ESCRT-0 in *Salmonella* pathogenesis, we investigated the impact of depeleting core and accessory ESCRT-0 host proteins on infection outcome. We employed a high-content single-cell microscopy screen in HeLa cells infected with either SL1344-WT or SL1344-Δ*sopB* (**Fig. 5a**). Cells were transfected with siRNA pools targeting candidate host proteins prior to infection and were imaged 6 hours post-infection. In parallel, we included cargo-selective retromer components VPS26A, VPS29 and VPS35 in the screen as these were enriched with SopB in Virotrap (**Fig. 1b**) and had been reported to positively impact intracellular *Salmonella* load upon knockdown [41]. We first quantified the percentage of infected cells under all experimental conditions. Under non-targeting control conditions, the average infection rates were 28.8 % for SL1344-WT and and 23.6 % for the Δ*sopB Salmonella* strain (**Fig. 5b**). Notably, depletion of the retromer subunits VPS26A, VPS29 and VPS35 significantly increased the proportion of infected cells, reaching 40.8 %, 52.0 % and 50.8 % for WT, and 33.8 %, 42.1 % and 49.1 % for Δ*sopB*, respectively. In contrast, depletion of ESCRT-0 components and accessory proteins had no significant impact on infection rates.

**Figure 5.**
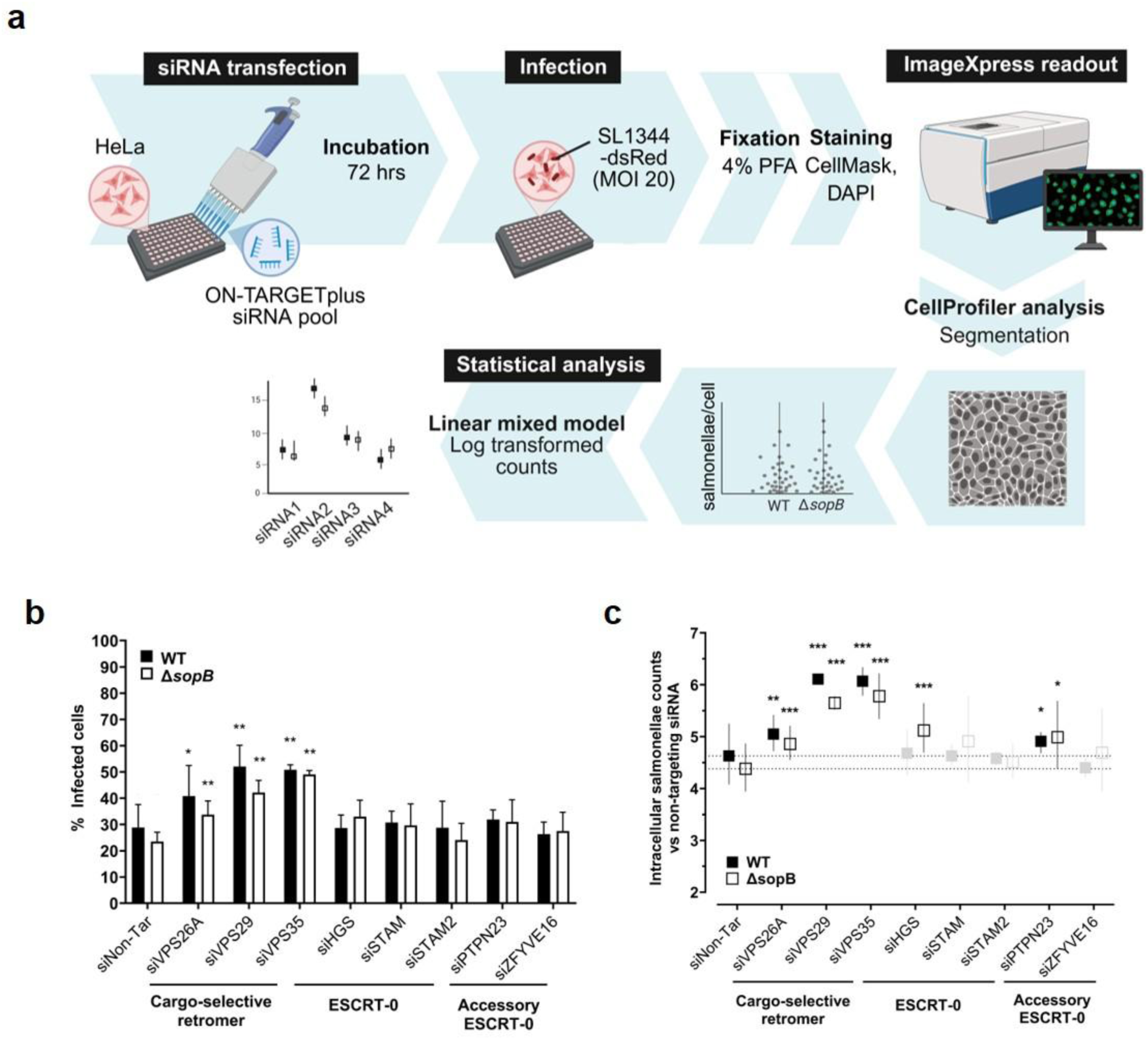
ESCRT-0 Modulates Salmonella Infection Burden in Coordination with SopB. (a) Schematic representation of the experimental setup employed for single-cell high-content imaging after bacterial infection upon knockdown of host factors. HeLa cells were treated with ON-TARGETplus siRNA pools against human host factors. After 72 hours silencing, cells were infected with either a SL1344-dsRed WT or ΔsopB strain at MOI 20 for 6 hours. Before readout, cells were fixed and the plasma membrane (CellMask) and nuclei (DAPI) were stained. Segmentation of the microscopy images was done in CellProfiler to quantify the amount of salmonellae per cell. These counts were log transformed and analysed using mixed linear model statistics. (b) Infected HeLa cells were quantified for both SL1344-WT and -ΔsopB strains separately. Cells were considered infected if at least 2 bacteria were present after 6 hours. Pairwise comparisons between each siRNA condition and the non-targeting control were performed separately for both strains. Data represents results from three independent replicate experiments (*p < 0.05, **p < 0.01). (c) Intracellular salmonellae per infected cells were quantified and pairwise comparisons between each siRNA condition and the non-targeting siRNA control were performed separately for both the WT and ΔsopB strains. Data represents results from three independent replicate experiments (*p < 0.05, **p < 0.01, ***p < 0.001). Data points in grey are not significant.

Taking advantage of the single-cell, and single-bacterium resolution of the screen, we next quantified the intracellular *Salmonella* burden per infected cell (**Fig. 5a**). We developed a statistical analysis pipeline to account for condition-specific variation in high-content screening data (see Materials and Methods). Comparison of WT and Δ*sopB* strains under control conditions revealed no significant difference in *Salmonella* burden. However, knockdown of each of the three retromer components led to a significant increase in intracellular *Salmonella* numbers per cell for both strains (**Fig. 5c**). Interestingly, HGS depletion significantly increased bacterial burden exclusively in Δ*sopB*-infected cells. Conversely, knockdown of the accessory protein HD-PTP (encoded by *PTPN23*) resulted in a mild but significant increase in bacterial burden, irrespective of Δ*sopB* background. These findings suggest that ESCRT-0 core component HGS restricts *Salmonella* burden in absence of SopB, pointing to a compensatory effect of SopB, whereas silencing of the accessory component HD-PTP increases bacterial burden regardless of SopB presence

## Discussion

*Salmonella* deploys a diverse arsenal of effectors to manipulate host cell signaling. Among these, the inositol phosphatase SopB, discovered over two decades ago [11], remains incompletely understood despite extensive study, with emerging functions continuing to reveal its multifunctionality [21, 23, 31, 32, 34, 42]. Dissecting the specific contribution of SopB requires precise mapping of its host interaction network. Using the sensitive Virotrap interactomics platform [24] (**Fig. 1a**), well-suited for proteins that are cytotoxic upon (over)expression [28], as is the case for SopB [11], we generated the most comprehensive SopB-host interactome to date that includes both novel and previously reported host interactions (**Fig. 1b, f-g**).

Virotrap captured known SopB-associated proteins, including the retromer subunit VPS35 [41] and multiple sorting nexins (SNX3, SNX9, SNX18) [13, 34, 42] (**Fig. 1b, h**). The complete recruitment of the cargo-selective retromer trimer (VPS26, VPS29, and VPS35), alongside SNX2 and SNX27—both known to localize to the SCV [43]—suggests extensive retromer engagement. SNX2 promotes SNX3-mediated tubule formation for mannose-6-phosphate receptor (M6PR) cycling [44, 45], a process known to be subverted by SopB [42], while SNX27 functions as a recycling adaptor directing cargo to the plasma membrane, rather than to SNX2 and SNX3, which route cargo to the trans-Golgi network (TGN) [46]. In addition, SNX9 and SNX18 that mediate plasma membrane remodeling independent of retromer [47], were also identified as potential SopB interactors (**Fig. 1b, h**). The diversity of SNX subfamilies recovered supports a model in which Virotrap captures the SopB interactome at both the plasma membrane and endosomal compartments, reflecting its multi-compartment localization at the invasion sites and the SCV [18, 19].

Moreover, our interactomics data position SopB within a broad host membrane–remodeling network, directly linking its phosphoinositide phosphatase activity to retromer, SNAP receptor (SNARE), and exocyst complexes (**Fig. 1f, g**). The enrichment of STX3, STX7, VAMP3, and VAMP7 with wild-type SopB compared to the catalytic SopB mutants suggest a role in selective fusion events involving host endo-/lysosomal compartments. Enrichment of EXOC1, -4 implies involvement with the complete heterotetrameric exocyst subcomplex. This aligns with recent evidence for SopB-dependent assembly of EXOC2 and EXOC3 at *Samonella* invasion sites [33]. Based on our data, SopB also engages SNX3, SNX9, and SNX18 through phosphoinositide remodeling, which is consistent with prior work [13, 34, 42]. Remarkably, the endosomal complexes enriched with SopB serve different modes of membrane remodeling: protein removal via internalization (ESCRT), tubular extrusion (retromer), fusion (SNARE), and membrane microdomain organization (clathrin and flotillin) that coordinates these events [48, 49]. Together, these results indicate that SopB mobilizes multiple endosomal machineries to modulate the membrane composition and dynamics of the SCV.

Among the novel insights revealed by the SopB Virotrap interactome, we identified a notableassociation between the effector and the complete host ESCRT-0 complex, comprising HGS, STAM and STAM2 (**Fig. 1a**), notably independent of SopB’s enzymatic activity (**Fig. 1h, j**). The ESCRT-0 machinery is a critical regulator of endosomal sorting, responsible for directing ubiquitinated receptors into the lumen of multivesicular bodies (MVBs) via ILV formation [26]. Upon activation, many cell surface receptors undergo ubiquitination and are internalized into early endosomes, where they are either recycled to the plasma membrane or targeted for lysosomal degradation. Through the sequential action of the ESCRT complexes (*i.e*., ESCRT-0, -I, -II, and -III), ubiquitinated cargo is sorted into ILVs, after which the MVB fuses either with the lysosome for receptor turnover or with the plasma membrane for exocytosis [50]. Next to receptor trafficking, ESCRT functions in processes such as micro-autophagy, autophagosome closure, membrane repair and cytokinesis [51], and is highly conserved across eukaryotes [52]. The phosphatase-independent enrichment of ESCRT-0 with SopB, together with the pivotal roles of both SopB and ESCRT in endomembrane dynamics, led us to explore this intriguing interaction further. Using NanoBiT assays, we confirmed direct binding between SopB and ESCRT-0 subunits HGS, STAM, and STAM2 (**Fig. 2c**). Additionally, confocal microscopy showed co-localization of HGS with heterologously expressed SopB at host endomembranes (**Fig. 2d-f**), consistent with the known localization of both SopB and ESCRT-0to PI(3)P-enriched membranes [20, 53]. Notably, HGS localization to macropinosomes appeared to be independent of SopB, as similar localization was observed upon SopE2 expression (**Fig. 2d**), another *Salmonella* T3SS-1 effector involved in membrane ruffling, bacterial entry and macropinosome formation [12, 54]. Considering ESCRT-0’s established role in cargo sorting within specialized membrane domains [49], SopB and HGS may be co-sorted into ILVs at these sites, potentially along with other host proteins. This hypothesis is supported by the reported presence of SopB on exosomes derived from *Salmonella*-infected cells [55]. To explore whether our observations are recapitulated during infection, we examined ESCRT-0 dynamics in infected cells. HGS, the core ESCRT-0 subunit, was observed to accumulate at IAMs shortly after bacterial entry and at SCVs approximately 30 minutes post-infection (**Fig. 3a-d)**, paralleling the established spatiotemporal distribution of SopB [56]. Notably, SopB enhances the recruitment of HGS to the SCV (**Fig. 3c**), suggesting it modulates ESCRT-0 localization during infection.

SopB is mono-ubiquitinated on multiple lysine residues in host cells [18, 19]. Mining our Virotrap data for diGly modifications corroborated reported ubiquitination sites at K19, K41, K93, and K541 (**Fig. 4a**). Ubiquitination has been associated with SopB relocation from the plasma membrane to the SCV in Caco-2 cells [19] and with prolonged SopB retention at the SCV in HeLa cells [18]. These lysines are notably conserved across diverse *Salmonella* serovars [18], underscoring their importance in *Salmonella* pathogenesis. Our NanoBiT assays further supported the role for ubiquitination in SopB - ESCRT-0 binding (**Fig. 4d**). Deletion of both HGS ubiquitin-binding domains (VHS and UIM) abolished this interaction. Recent research suggests that multimeric ESCRT-0 complexes assemble in response to localized increases of PI3P and achieve high-avidity binding via multiple low-affinity ubiquitin contacts [53, 57]. Consequently, disrupting both ubiquitin-binding domains of HGS is likely necessary to disrupt SopB-ESCRT-0 binding.

Functional interrogation via siRNA-mediated depletion of core and accessory ESCRT-0 components during infection with WT and Δ*sopB* SL1344 revealed that loss of these host proteins did not impact the proportion of infected cells (**Fig. 5b**), suggesting that ESCRT-0 is not involved in bacterial entry, unlike its previously reported role in viral entry [58]. Strikingly, depletion of HGS differentially influenced intracellular bacterial loads depending on the presence of SopB (**Fig. 5b-c**), pointing to a multilayer functional interplay between this host factor and *Salmonella* proteins. Especially, bacterial burden increased upon HGS silencing only in the absence of SopB (**Fig. 5c**), suggesting that SopB may counteract an anti-replicate effect of HGS. Given the established role of SopB in modulating phospholipid composition and protein recruitment at the SCV [12, 13], we propose a model in which SopB initially promotes HGS recruitment at the SCV (**Fig. 3c**, **Fig. 4e**), and subsequently interferes with its function by engaging its ubiquitin-binding domains.

The evolutionary conservation and central role of the ESCRT machinery make it an attractive target for intracellular pathogens, yet direct exploitation by *Salmonella* effectors has not been previously reported. Our findings establish SopB as a bacterial effector that engages and recruits the host ESCRT-0 complex. The distinctive mono-ubiquitination of SopB, together with its ubiquitin-dependent interaction with HGS, supports a model in which SopB acts as a molecular mimic of ubiquitinated host cargo, possibly modulating SCV membrane composition (**Fig. 4e**). Whether such sophisticated mimicry extends to other pathogens remains an open and compelling avenue for future research.

## Acknowledgments

We want to thank Eef Parthoens and colleagues of the VIB BioImaging Core Ghent and Rudmer Postma at LUMC for their valuable contribution to the research presented in this paper.

## Contributions

Margaux De Meyer: conceptualization, formal analysis, investigation, methodology, visualization, writing – original draft, writing – review & editing. Annick Verhee: investigation. Hanna Grzesik: investigation. Delphine De Sutter: investigation. Jon Huyghe: methodology, resources. Louis Delhaye: investigation. Tessa Van de Steene: investigation. Igor Fijalkowski: formal analysis, software. Veronique Jonckheere: investigation. Leander Meuris: formal analysis, software. Mathieu JM Bertrand: methodology, funding acquisition. Petra Van Damme: conceptualization, methodology, funding acquisition, writing – review & editing. Virginie Stévenin: conceptualization, investigation, formal analysis, methodology, resources, supervision, funding acquisition, writing – review & editing. Sven Eyckerman: conceptualization, methodology, supervision, funding acquisition, writing – review & editing.

## Materials and Methods

### Cell Culture

Authenticated HEK293T cells originate from [59] (Rufer lab, CHUV, Lausanne) and were maintained in high-glucose Dulbecco’s Modified Eagle Medium containing GlutaMAX (DMEM; Gibco, cat no. 10566016), supplemented with 10% fetal bovine serum (FBS, Gibco, cat no. 10270106) and 100 U/mL penicillin/streptomycin (Gibco, cat no. 15070063). HeLa cells were cultured in Minimum Essential Medium (MEM; Gibco, cat. no. 11534496), supplemented with non-essential amino acids (Gibco, cat. no. 11140035) and sodium pyruvate (Gibco, cat. no. 11360070). Cells were kept in a humidified incubator at 37 °C and 5% CO2.

### Cloning and Plasmids

Cloning was performed using *Escherichia coli* strain DH10B with standard chemical transformation. *Salmonella* Typhimurium SL1344 genomic DNA was extracted as described in [60] and served as a template for *sopB* coding sequence amplification. Human coding sequences were PCR-amplified from cDNA synthesized using the SuperScript IV first-strand synthesis system (Thermo Fisher, cat. no. 18091050). The cDNA was derived from RNA isolated from HEK293T or HeLa cells using TRIzol reagent (Thermo Fisher, cat. no. 15596026). Amplicons were purified by means of a Nucleospin® Gel and PCR clean-up kit (Machinery-Nagel) or E.Z.N.A.® Plasmid DNA Mini Kit II. (Omega Biotek) according to the manufacturers’ instruction. Coding sequences were cloned into the Gateway® pDONR221 and pMET7-GAG-SP1 [24] using BP Clonase II Enzyme (Invitrogen) and LR Clonase II Plus Enzyme (Invitrogen), respectively, for Virotrap. The pMD2.G (VSV-G envelope-expressing plasmid; Addgene, plasmid no. 12259) was retrieved from Addgene, the pcDNA3-FLAG-VSV-G (Addgene, plasmid no. 80606) plasmid was provided by the Eyckerman lab and the pSVsport vector was obtained from Life Technologies. Other constructs were created using the BsaI-based Golden Gate assembly cloning system as described in [61] for genetic fusions with EGFP, mScarlet, LgBiT-FLAG and SmBiT-V5. Mutant SopB constructs were generated from the wild-type SopB coding sequence using the Quickchange II site-directed mutagenesis kit (Agilent, cat. no. 200523).

### Stable Cell Line Generation

To generate HeLa cell lines stably expressing HGS-mScarlet, lentiviral constructs were prepared encoding HGS-mScarlet along with a puromycin resistance cassette for antibiotic selection. Subsequently, these constructs were used to produce lentiviruses following the procedure outlined in [61]. HeLa cells equipped with the doxycycline-inducible rtTA/tTS system, as detailed [62], were transduced with HGS-mScarlet lentivirus at an MOI of 1. The medium was replaced the next day. Two days after transduction, 1 µg/mL of puromycin was introduced into the cell culture medium, based on previously established kill-curves in the lab. Selection using puromycin continued for at least two days, or until no cells from the parental lines survived under the same conditions.

### Virotrap

Virotrap was performed as described previously [63, 64]. In brief, ten million HEK293T cells were seeded per T75 flask in complete DMEM and transfected the next day using polyethylenimine (PEI) reagent (linear 25 kDa, Polysciences, Inc.). The transfected DNA/PEI mixture consisted of 0.71 μg pcDNA3-FLAG-VSV-G, 0.36 μg pMD2.G, 6.43 μg pMET7-GAG-SP1-sopB (mutant) or 3.75 µg pMET7-GAG-SP1-eDHFR and 2.67 µg pSVsport (control samples) and 37.5 μL PEI (1 mg/mL solution in MQ, pH 7.0) per T75. The cellular supernatant was harvested 48 h after transfection, spun at 1500xg for 3 minutes at room temperature and filtered through a 0.45 μm Millex® filter (Millipore). Per sample (equivalent of T75), 20 μL MyOne Streptavidin T1 beads (10 mg/mL; Invitrogen), washed in 20 mM TRIS HCl pH 7.5 and 150 mM NaCl, was loaded with 2 μL anti-FLAG BioM2-biotin antibodies (1 mg/mL; ANTI-FLAG® BioM2, cat. no. F9291, Sigma Aldrich) in 200 μL washing buffer by end-over-end rotation and incubation for 2 h. VLPs were allowed to bind the anti-FLAG-coated beads for 2 h by end-over-end rotation at room temperature. Bead-bound VLP complexes were washed once with washing buffer and eluted using 20 µL elution buffer (20 mM TRIS HCl pH 7.5, 150 mM NaCl, 200 µg/ml FLAG-peptide) and incubate for 30 minutes at 37°C. Subsequently, VLPs were lysed by addition of 2.2 µL amphipathic polymer solution (Amphipol A8-35, Anatrace; final concentration of 1 mg/mL) and incubation for 10 minutes. For protein concentration, proteins were pelleted from the lysates by acidification (0.2% final concentration formic acid) [65]. After acidification, protein pellets were dissolved in 20 μL of 50 mM triethylammonium bicarbonate (TEAB) buffer (pH 8.5), boiled and digested overnight using 0.5 μg of sequence-grade modified trypsin (Promega). After a final acidification step (0.4% formic acid, final concentration), samples were separated on an UltiMate™ 3000 RSLCnano (Thermo Scientific) and analyzed on a Q Exactive HF instrument (Thermo Scientific; 7.5 μL injected, 1.5 h long run) as described previously [66, 67].

### Mass spectrometry Data Analysis

Searches were performed using MaxQuant (Version 1.6.6.0) [68] against the human SwissProt Proteome Database (Release 2022-02) complemented with eDHFR, FLAG-VSV-G, VSV-G, Gag and SL1344 SopB protein sequences. In MaxQuant, multiplicity was set to one, indicating that no labels were used. Furthermore, we performed label-free quantification (LFQ) using MaxQuant’s standard settings with a minimum of two ratio counts and considering unique and razor peptides for protein quantification. A decoy database of reversed protein sequences was used to estimate FDR, and 1% FDR threshold was applied. Di-glycine, methionine oxidation, and N-terminal protein acetylation were set as variable modifications and trypsin/P was set as the digestion enzyme allowing two missed cleavages. N-terminal acetylation was included in protein quantification. The MaxQuant ProteinGroups data file was processed using R Studio (R Foundation for Statistical Computing, V1.3.959) and custom R scripts. The dataset was further filtered based on reversed hits, potential contaminants and proteins only identified by site. For identifications with LFQ values calculated for all bait replicates (*i.e*., 4 valid values), LFQ intensities were log2 transformed and missing values were imputed using the QRILC function with default parameters from the imputeLCMD package in R [69]. Significance was assessed using limma in R [70] as previously demonstrated [60]. Replicate samples were grouped and compared to the control samples (Gag-eDHFR or Gag-SopB-WT) in a pairwise analysis. Basic data handling and Pearson correlation calculations were performed in Perseus V1.6.6.0 [71]. Network analysis was done using the STRING database [72] and Cytoscape [73] for additional visualization and ClueGo [74] enrichment analyses.

### Infection

*Salmonella* infections were conducted following the procedures described in [75]. Bacteria were cultured overnight in 3 mL of LB containing 0.3 M NaCl and 50 μg/mL of either ampicillin or kanamycin at 37°C shaking at 220 r.p.m. On the day of infection, the bacteria were further cultured at a 1:21 dilution in the same medium and conditions for 3 hours until the late exponential growth phase. Following this, the bacteria were washed and resuspended in EM buffer (120 mM NaCl, 7 mM KCl, 1.8 mM CaCl_2_, 0.8 mM MgCl_2_, 5 mM glucose, 25 mM HEPES, pH 7.3). The bacterial density was assessed using optical density at 600 nm and adjusted to the required multiplicity of infection (MOI) in warm EM buffer. All infections were performed at an MOI of 20. Cells were rinsed with EM buffer before bacterial addition and incubated for 30 minutes at 37°C in 5% CO_2_. Subsequent to this, cells were washed three times with sterile EM buffer to remove any extracellular bacteria. Depending on the specific experimental timeline, cells were either immediately fixed or continued to be incubated in EM buffer containing 10% FBS and 100 μg/mL gentamicin for one hour. For extended time points, the medium was replaced after one hour with EM buffer containing 10% FBS and gentamicin at 10 μg/mL.

### Confocal Microscopy

HeLa cells were seeded into each well of an 8-well imaging μ-Slide (Ibidi GmbH, cat. no. 80826). For transient expression studies, cells in each well were transfected with 100 ng of DNA using a 1:3.5 ratio of Fugene HD (Promega, cat. no. E2311) for 18 hours before imaging. Transfection mixtures were prepared in reduced serum Opti-MEM medium (Gibco, cat. no. 15392402). Induction of HeLa-HGS-mScarlet cells was done using 20 ng/mL doxycycline for 18 hours. For fixed samples, cells were treated with 4% PFA in PBS, which was pre-warmed to 37°C, for 10 minutes at room temperature (RT). After fixation, cells were washed three times for 5 minutes in PBS and were stained using DAPI (Thermo Fisher Scientific, cat. no. D1306) at a dilution of 1:1,000 in PBS for 15 minutes at RT. After three washes using PBS, cells were preserved in PBS at 4 °C. Microscopy acquisitions were performed on either an Andor Dragonfly 200 (Oxford Instruments) spinning disk microscope equipped with a 63x/1.40-0.60 oil Plan Apo objective, and an Andor Zyla 4.2 PLUS sCMOS Camera (Fig. 3; S2), or LSM880 Airyscan (Carl Zeiss, Jena, Germany) equipped with either a Plan-Apochromat 63x/1.4 Oil, DIC M27 (Fig. 2). The LSM880 utilized ZEN Black 2.3 SP1 software with lasers of 405, 561, and 633 nm, and the 488 nm line from an argon laser. The corresponding filter sets included BP 420-480, LP570, LP645, and BP 495-550, applied with the Airyscan detector in SuperResolution (SR) mode or in full Airyscan. For live-cell imaging, fast Airyscan mode was used. Post-acquisition, images were processed for pixel reassignment and 2D Wiener deconvolution in ZEN Black. Co-localization quantification was performed using ZEISS arivis software.

### NanoBiT

HEK293T cells were cultured in black 96-well tissue culture plates (Greiner, cat. no. 3916) at a seeding density of 15,000 cells per well in complete DMEM. Cells were incubated at 37°C with 5% CO_2_ for 16 hours to allow for attachment. Triplicate were prepared in 96-well plates. For each construct, 40 ng plasmid DNA was mixed with a final concentration of 0.25 M CaCl_2_ in sterile water per well to be transfected. This mixture was combined with an equal volume of 2X HeBs (Sigma Aldrich, cat. no. 51558), incubated for 10 minutes at room temperature, and vortexed before transferring the combined transfection mixture to the well. After incubating for 20 hours at 37°C, the Nano-Glo® Live Cell Substrate (Promega, cat. no. N2012) was mixed according to the manufacturer’s instructions and added to cells by replacing the culture medium with the substrate mixture. Plates were gently swirled to distribute the substrate, and luminescence measurements were performed at 37°C using EnVision (PerkinElmer, 2102 Multilabel reader). Cells were washed using PBS and frozen at -20 °C for subsequent lysate preparation.

### SDS-PAGE and Western Blotting

VLP and producer cell lysates were supplemented with sample loading buffer (XT sample buffer, Bio-Rad) and reducing agent (XT reducing agent, Bio-Rad) according to the manufacturer’s instructions. Proteins were separated on 4–12% gradient XT precast Criterion gel (Bio-Rad, cat. no. 3450123) using XT-MOPS buffer (Bio-Rad) at 150 V and subsequently transferred onto a PVDF membrane. For NanoBiT expression analysis, cells were lysed in 20 µL LDS loading buffer (4X, diluted in TBS, Genscript, cat. no. M00676) per well. Prior to separation on a 4–20% polyacrylamide gradient gel (GenScript, cat. no. M42015), protein samples were boiled for 10 minutes at 98 °C. Next, the protein samples were transferred onto a PVDF membrane for 3 hours at 60 V in blotting buffer (48 mM Tris, 39 mM glycine, 0.0375 % SDS and 20 % methanol). Membranes were blocked for 30 min in 1:1 Tris-buffered saline (TBS)/Odyssey blocking solution (cat no. 927-40003, Li-COR) and probed using primary antibodies in TBS-T/Odyssey blocking buffer. After three washes of 10 min in TBS-T (0.1% Tween-20), membranes were incubated with secondary antibody (1/5000 dilution; IRDye antibodies; Li-COR) for 30 min in TBS-T/Odyssey blocking buffer. Following three washes in TBS-T and one additional wash in TBS, fluorescent detection was done using an Odyssey infrared Fc imaging system (Li-COR) or Odyssey infrared scanner (Li-COR).

### High Content Microscopy Screen

In each well of a PhenoPlate 96-well plate (PerkinElmer, cat no. 6055300), 6000 HeLa cells (Neefjes Lab, endogenously expressing LAMP1-eGFP) were seeded and cultured in DMEM containing GlutaMAX (Gibco, cat. no. 10566016), supplemented with 10% heat-inactivated FBS. Following an overnight incubation, cells were transfected with either 20 nM siRNA or a non-targeting pool (Dharmacon, ON-TARGETplus SMARTpool) using DharmaFECT 1 transfection reagent (Dharmacon, cat. no. T-2001), in accordance with the manufacturer’s instructions. Subsequently, 72 hours after transfection, cells were infected with dsRed-expressing salmonellae at an MOI of 20. Fixed cells were obtained by treating with 4% paraformaldehyde (PFA) in PBS and subsequently stained with DAPI (1/500 in PBS) and Cell Mask Plasma Membrane staining, Deep Red (Thermo Fisher Scientific, cat. no. C10046, 1/500 in PBS). Imaging was performed at 40X magnification on an ImageXpress Micro confocal high-content instrument, capturing 16 fields per well. A Z-stack of 16 images with a step size of 0.5 µm was acquired. Z-projections were analyzed using CellProfiler [76], essentially as described by Voznica and colleagues [77]. Significant changes in percentage infected cells and mean salmonellae counts were determined using a custom R script. Briefly, bacterial counts were log-transformed to normalize the right-skewed distribution typical of count data, and condition-specific means were calculated. Cells containing at least 2 bacteria were classified as infected, while those with ≥30 bacteria per cell were treated as outliers and excluded from the analysis. Log-transformed means were analyzed using linear mixed models with random intercepts for both experiment and block.

Supplementary Video 1. Cropped view of time-lapse microscopy of HeLa cells expressing mScarlet-HGS upon doxycycline induction (white) and infected with GFP-expressing *Salmonella* SL1344 WT (green). Bacteria are added to the cells just before the beginning of the acquisition to visualize entry and early events.

Supplementary Video 2. Cropped view of time-lapse microscopy of HeLa cells expressing mScarlet-HGS upon doxycycline induction (white) and infected with GFP-expressing *Salmonella* SL1344 WT (green) in the presence of fluorescent Dextran (yellow). Dextran and bacteria are incubated with HeLa cells for 30 min and washed (W) before the beginning of the acquisition to visualize IAM dynamics.

---

**Supplementary Figure 1.**
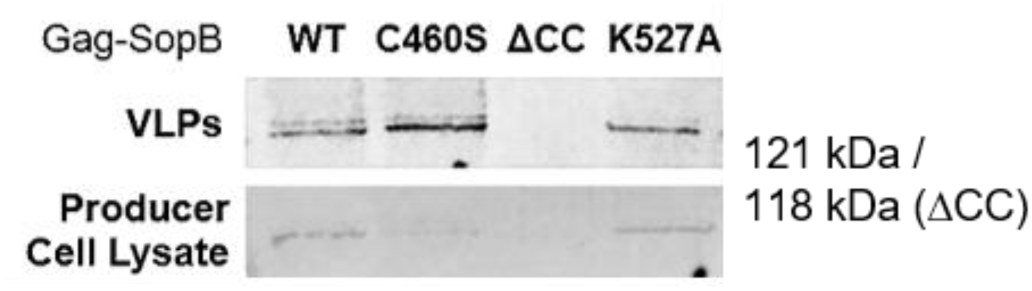
Immunoblot analysis of Gag-SopB virus-like particles (VLPs). VLPs from HEK293T cells expressing Gag-SopB constructs were captured from culture supernatants by anti-FLAG immunoprecipitation. Wild-type (WT), catalytic mutant (C460S), coiled-coil deletion mutant (omitted from our study since no expression was obtained), and reduced catalytic mutant (K527A).

**Supplementary Figure 2.**
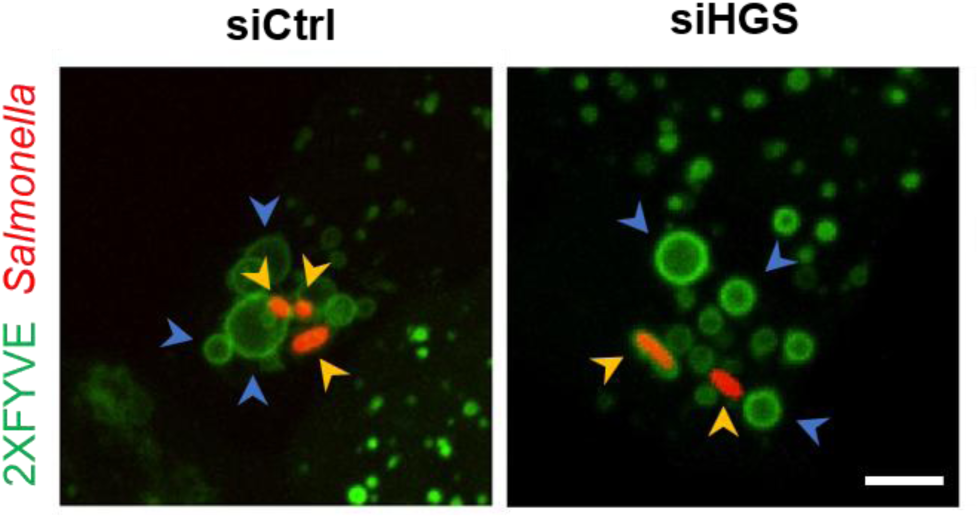
IAM and SCV formation during SL1344 invasion upon HGS depletion. HeLa cells were subjected to HGS depletion using siRNA for 72 hours and were infected with SL1344-WT(dsRed). All images display infection sites within a few minutes after bacterial entry (i.e., timeframe following ruffle formation). Yellow arrowhead: SCV, Blue arrowhead: IAM. Scale bar indicates 5 µm.

